# Data and Diversity Driven Development of a Shotgun Crystallisation Screen using the Protein Data Bank

**DOI:** 10.1101/2021.08.11.456002

**Authors:** Gabriel Abrahams, Janet Newman

## Abstract

Protein crystallisation has for decades been a critical and restrictive step in macro-molecular structure determination via X-ray diffraction. Crystallisation typically involves a multi-stage exploration of the available chemical space, beginning with an initial sampling (screening) followed by iterative refinement (optimisation). Effective screening is important for reducing the number of optimisation rounds required, reducing the cost and time required to determine a structure. Here, we propose an initial screen (Shotgun II) derived from analysis of the up-to-date Protein Data Bank (PDB) and compare it with the previously derived (2014) Shotgun I screen. In an update to that analysis, we clarify that the Shotgun approach entails finding the crystallisation conditions which cover the most diverse space of proteins by sequence found in the PDB - which can be mapped to the well known Maximum Coverage problem in computer science. With this realisation we are able to apply a more effective algorithm for selecting conditions, such that the Shotgun II screen outperforms the Shotgun I screen both in protein coverage and quantity of data input. Our data demonstrates that the Shotgun I screen, compared with alternatives, has been remarkably successful over the seven years it has been in use, indicating that Shotgun II is likely to be a highly effective screen.

## 1. Introduction

Macromolecular structure determination through the technique of X-ray diffraction has been wildly successful. A public repository of protein structures was initiated in 1971 (Bernstein *et al*., 1997). Fifty years later there are close to 175,000 structures in the Protein Data Bank (PDB, (Burley *et al*., 2017)), of which over 150,000 have been determined using X-ray crystallography. The bottleneck of the X-ray diffraction technique is widely appreciated to be the production of appropriate crystals. Without this constraint it is not clear how many crystal structures would be available today. The ‘crystal bottleneck’ is really a misnomer – problems with protein production, crystal production and diffraction quality are generally lumped together under the umbrella phrase “couldn’t get crystals”.

Given sample, the process of generating a suitable crystal for diffraction studies requires an initial search of crystallisation space (screening), followed - almost certainly - by one or more cycles of refinement of any hits found in the initial search (optimisation) (Bergfors, 2009; McPherson & Gavira, 2013; Luft *et al*., 2014). The chemical space that needs to be searched is vast - a rough estimate suggests about 20 billion possibilities^**1**^. Experience tells us that crystallisation chemical space is not uniformly successful - certainly more proteins have been crystallised from conditions at pH 7 than at pH 3 (Fig. B.1). The successful crystallisation landscape for any one protein might look quite different from the successful crystallisation space for another protein. However, if we have information about the successful crystallisation spaces for many different proteins, we might be able to find general ‘hotspots’ in crystallisation chemical space, where many different proteins could form crystals (Fig. 1).

**Figure 1.**
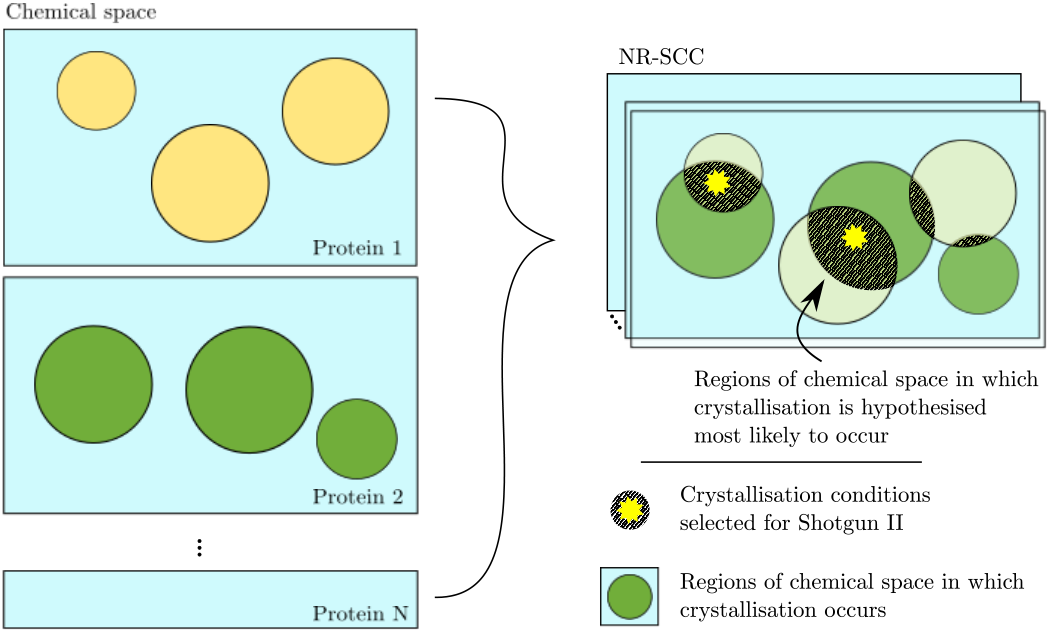
Crystallisation chemical space, represented here by blue rectangles, is a large space. Every (crystallisable) protein will have one or more success regions within that space that supports the formation of crystals - these are represented by coloured circles within the blue rectangles. If many proteins crystallise in the same part of crystallisation space that would be a general crystallisation ‘hotspot’, and might be a good place to start a new crystallisation campaign. The current study tries to find the overlap between two diverse spaces - protein space, and chemical space. These diagrams are cartoons: the dark hotspots are showing overlap from only from two proteins, and the success regions are probably asymmetrical and of course much smaller than the whole of chemical space.

When starting a crystallisation campaign with a novel protein, testing the successful regions of chemical space or hotspots would be a sensible place to start. Commercially available screens (which are assumed to cover the hotspots in crystallisation chemical space) are almost invariably used for the initial screening of crystallisation space. The first crystallisation screen was described by Jancarik and Kim in 1991 (Jancarik & Kim, 1991). That paper presented a set of 50 crystallisation conditions and showed that this small number of conditions could be used to find initial crystal hits from proteins that had already been crystallised and new proteins for which crystallisation conditions were not known. The conditions were created by combining a pH buffer, an additive and a precipitant. The buffers, additives and precipitants were chosen by ‘trial and error’ from a list of chemicals which were known to have been used successfully in previous crystallisation trials. The screen contained conditions which were later shown to be successful, rather than consisting of complete conditions that had already been successful in crystallising proteins. This 50-condition screen was commercialised later in 1991 (and is still available: Crystal Screen, from Hampton Research, https://hamptonresearch.com/product-Crystal-Screen-Crystal-Screen-2-Crystal-Screen-HT-1.html). Since then, a number of companies have provided pre-mixed crystallisation screens. Some of these commercial screens are based on successful conditions - either from a single laboratory (*e*.*g*. the JCSG screens or the MCSG screens) or from data mining the PDB, (*e*.*g*. the MemGold screens). Not all of the commercial screens are unique; most vendors include a screen which - like the Hampton Research Crystal Screen - is based on the original ‘sparse matrix’ screen from Jancarik and Kim. The C6 webtool (c6.csiro.au, (Newman *et al*., 2010)) shows that there are over 22,000 commercial crystallisation conditions available for purchase, but only 12,000 of these are different from each other (*i*.*e*. distinct). One of the perennial issues in crystallisation is deciding which screening kit(s) to use in the initial stages of a crystallisation campaign. It would be incredibly useful if there was validated experimental data available about which commercial kits are most successful, and even better if there was solid data about which different kits are more or less well suited for different types of samples. To determine the success of a screen, we would need to compare how many positive outcomes (crystalline hits) are obtained for a given number of setups. Ideally, many different diverse proteins would have been used to test the kit - or in the case of directed initial screens, many different instances of the class of protein being targeted. Unfortunately, both the numerator (number of hits) and the denominator (number of setups) data are almost impossible to obtain. One could perhaps estimate the number of times a screen has been set up by looking at the number of kits that have been purchased. However, while the sales numbers are known to the vendor, they might be reluctant to share them. More importantly, just because a screen was purchased does not necessarily mean that it was used to set up experiments. The outcomes of any screening of commercial conditions are even less easy to estimate. Firstly, there is no clear definition or indeed understanding of what a ‘crystalline hit’ is within the structural biology community. Secondly, there is no reliable method of communicating the complete outcome of a crystallisation campaign. If the campaign were successful, and resulted in a crystal structure which was published or deposited in the PDB, only the information about the crystallisation condition used to grow the crystal used in the diffraction experiment would have been recorded. There is currently no standard mechanism for capturing information about any other crystal hit(s) found during a crystallisation campaign.

In 2014, we performed an analysis which compared commercially available crystallisation conditions with successful conditions extracted from the Protein Data Bank (Fazio *et al*., 2014). The comparison was used to create a 96-condition screen of the observed most successful (commercial) crystallisation conditions. We were in essence trying to find which - if any - of the commercial conditions were associated with hotspots in general crystallisation space. We called this set of 96 conditions the ‘Shotgun’ screen (hereafter called Shotgun I), after the screening approach used by Page *et al* during the era of Structural Genomics (Page *et al*., 2003). Since then, the number of protein structures determined by X-ray methods and deposited in the PDB has increased by 50,000, and we wondered if these new depositions would refine our ability to locate general hotspots in crystallisation space.

The Shotgun I screen was created by parsing the REMARK280 (R280) of the PDB for crystallisation conditions then filtering by sequence, so that only distinct crystallisation conditions were returned for each sequence. This produced a list of ‘non-redundant successful crystallisation conditions’, or NR-SCC. We also filtered the list of commercially available conditions to produce a list of all the distinct crystallisation conditions available for purchase – the ‘distinct commercial crystallisation conditions’ or D-CCC. These two lists were then used to create a histogram of the intersection between the two lists, where the histogram was ordered by the number of times each condition shared between the two lists was found in the NR-SCC. The first 96 of these shared conditions – the most populated 96 conditions - made up the Shotgun I screen.

The Shotgun I screen has been in use in our laboratory (CSIRO’s Collaborative Crystallisation Centre - C3 - in Melbourne, Australia) since 2014 and we have in-house data from over 4,000 protein samples that have been set up using the screen. The in-house data includes scores associated with images of the drops set up with the Shotgun I screen. Thus we have, for our laboratory, reasonable estimates to enable us to calculate the success of the Shotgun I screen along with other screens used in our laboratory.

## 2. Methods

All data analysis was performed using Python with the SciPy(Virtanen *et al*., 2020) and pandas packages.

### 2.1. Parsing the PDB

*All code/scripts used to parse the R280 fields will be made available at https://github.com/gabi-a/Shotgun-II*

A crystallisation condition here is taken to mean only the chemical cocktail that is mixed with the protein sample to engender crystallisation, with no consideration being given to the type of experiment (*e*.*g*. vapour diffusion, microbatch), the physical parameters (*e*.*g*. temperature, drop ratios, drop size) or the protein formulation which are also part of the crystallisation experiment. A condition contains one or more chemical factors. Each factor consists of a chemical name, with an associated concentration, unit and possibly a pH. The free-text R280s need to be parsed to give a collection of complete and consistent descriptions of the crystallisation conditions associated with the PDB. For the current analysis we used the same Python scripts as were used in 2014. These scripts try to identify which part of the text might refer to the precipitant, then search for likely chemical names, and try to associate concentrations, units and potentially pH values to the identified chemicals. Once a putative chemical name is located, excess white space is removed, the case is rationalised and any waters of hydration are also removed. The next step standardises the chemical names using either an alias table or a substitution table. The alias table strictly resolves naming convention issues, *i*.*e*. all names which map to the same alias are the same chemical. ‘NaAc’, ‘Na Acetate’ and ‘NaOAc’ are all aliases of ‘sodium acetate’. There are some common abbreviations in the PDB which are ambiguous. For example ‘citrate’ usually means ‘trisodium citrate’ or some combination of the acid and base ‘trisodium citrate-citric acid’. We address this by applying substitutions to ambiguous names, where the replacement names are chosen by how many times they appear in the set of commercial screens. Note that substitutions are different to aliases: the aliases map identical chemicals to a single name, whereas the substitutions resolve ambiguities using an educated guess.

There is only one level of translation. If a complete description is given which does not use chemical names, concentrations and units, it will not be recognised. Thus ‘crystallised out of Index Screen A1’, despite being complete and unambiguous, would not be successfully parsed to (2 M ammonium sulfate, 0.1 M trisodium citrate-citric acid, pH 3.5). Similarly, we do not recognise screen specific mixture names - *e*.*g*. ‘Precipitant Mix 1’ from the Morpheus screen.

The pH of a condition is estimated by the average of the pH of each buffering factor in the condition. A factor is classified as a buffer if and only if the R280 has a pH and the pH is within 1 unit of the pKa: |pH − pKa| ≤ 1.

To compare chemical concentrations, we convert all units (M, w/v, v/v) to w/v using the best information available. In the few cases where density of a chemical is not known we assume a density of unity. If the solubility of a chemical is known and the parsed concentration exceeds that concentration, we set the concentration to ‘unknown’ and do not use it in further analyses.

To estimate the efficiency of translation, we selected two random sets of 100 R280s, and compared the original text to the parsed condition by hand. We did not limit the selection to conditions deposited since the 2014 work.

The ion concentrations used in the C6 comparison are determined by further parsing of the chemical names, taking stoichiometry into account - *e*.*g*. ‘0.2 M trisodium citrate’ is parsed as ‘3×0.2 M=0.6 M sodium, 0.2 M citrate’.

### 2.2. NR-SCC

Sequence clusters at 95% identity using MMSeqs2 were downloaded from https://www.rcsb.org/docs/programmatic-access/file-download-services. The downloaded output was clustered by chain identity. The data were re-clustered so that each cluster consists of PDB entries which all have the same chains, forming a distinct set of unique combinations of chains. The distinct set of parsed conditions associated with each cluster constitutes the NR-SCC.

### 2.3. D-CCC

The alias and substitution tables used to parse the PDB were also used to create a set of commercial conditions with consistent naming and units. Removing any duplicate conditions from the commercial screens gives 12,484 conditions, these make up the set of Distinct Commercial Crystallisation Conditions (D-CCC). Many of the conditions in the D-CCC are found in more than one screen. For these conditions we simply don’t know which screen was set up, although we can narrow down the possibilities. The most abundant condition in the PDB data (0.2 M magnesium chloride, 30 % w/v polyethylene glycol 4000, 0.1 M tris pH 8.5) is found 30 times in various screens available from 8 different vendors.

### 2.4. Coding and Algorithms

A number of tables (pandas dataframes) were used to store the data for analysis, but converted to NumPy or standard python objects for processing. Each chemical factor from the PDB and commercial screens was stored in a table with a unique ID and with columns: [name, concentration (parsed), units (parsed), pH (parsed), percentage concentration (derived), molarity (derived), ions (derived), buffer]. The NR-SCC and D-CCC are then simple python objects (list, dict) on which data analysis is fast, with entries being the IDs of the factors, *e*.*g*. a condition is a tuple of factor IDs sorted lowest to highest such that it can be used as a key for fast searching. A linking table between NR-SCC and D-CCC conditions was stored as a pandas dataframe and converted to two NumPy arrays (NR-SCC to D-CCC and vice-versa) for fast lookup.

### 2.5. Maximum Coverage Problem

The maximum coverage problem is solved by implementing the greedy algorithm with uniform weights (Chvatal, 1979). The 96 conditions are selected by iteration. Initially, the screen is empty and the possible conditions are all those found in the NR-SCC. At each step the condition which would result in the largest increase in the screen’s coverage of protein space is added to the screen and removed from the available choices. For a 96 condition screen, 96 iterations are required, however in order to refine the screen using chemical diversity (Section 3.6.2) roughly 400 iterations were required.

### 2.6. C6 Metric

To calculate the internal diversity, or the average distance between any two of the conditions within a screen of a screen, we use the C6 distance metric (Newman, 2011), which provides a heuristic for estimating the similarity between two crystallisation conditions. The C6 metric is the Manhattan distance between two condition vectors where each term is normalised by solubility, either using the maximum or average maximum concentration over commercial space (the D-SCC) of the species being compared. There are four components of condition space accounted for: factor concentrations, ion concentrations, PEG concentrations and pH. For the purposes of this work, the original C6 metric was reimplemented in Python using the following relationships:

The metric is a “dissimilarity” metric; it is 0 if the conditions are identical and 1 if they have no similarity. The distance between two conditions *i* and *j* is given by:

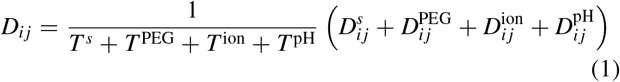

Where *T*^*s*^ is the number of distinct factors, *T*^PEG^ are the number of distinct PEGs with MW within a factor 2 of each other, *T*^ion^ is the number of distinct ions and *T*^pH^ is 1 if both conditions have an associated pH and 0 otherwise (pH may not be available due to parsing or lack of available data). Each component *D* combined with it’s associated pre-factor *T* may be thought of as “pulling” the metric closer to 0.

The factor term evaluates the distance between factor concentrations normalised by solubility:

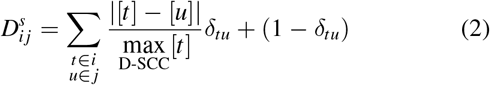

Where *δ* is the Kronecker-delta, [t] is the concentration of factor *t*, and we implicitly do not double count.

The PEG term is included so that PEGs which are within a factor 2 molecular weight are seen as similar:

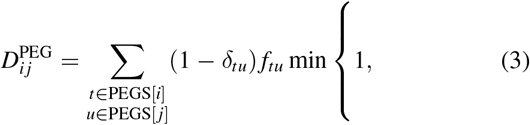

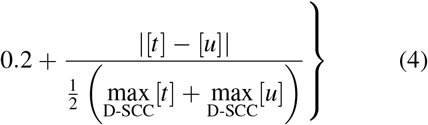

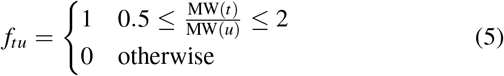

The ion term means that conditions containing non-identical factors which share the same ions are seen as similar:

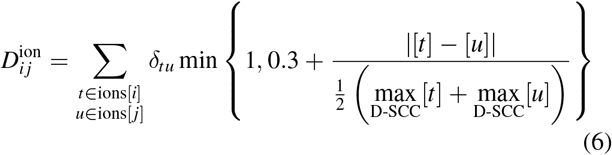

Finally the pH term is the distance between each conditions buffered pH:

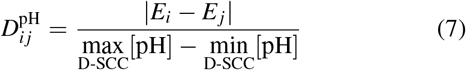

where *E*_*i*_ is the estimated pH of condition *i*.

### 2.7. Co-hits

A ‘hit’ refers to a condition crystallising a specific protein. Two conditions ‘co-hit’ if they both tend to crystallise the same proteins. We define co-hit to be the probability that A and B Hit given A or B Hit:

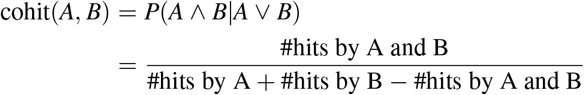

To compare the C6 score and co-hits we calculate each score for every pair of conditions within a screen (primarily the Shotgun I screen) - see Fig. 3 (a). The correlation coefficient *r* is calculated using the formula:

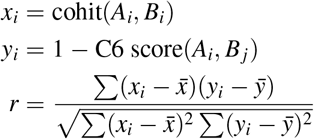

## 3. Results and Discussion

The Shotgun I screen has been set up over 4000 times in our crystallisation facility, which allows us to estimate the effectiveness of Shotgun I and to compare it to other initial screens available there.

### 3.1. Estimating the fitness of the Shotgun I screen

To estimate the success rate of any screen, we need to know both the number of times it produced a hit, and the total number of times it was set up. To compare the success rate of different screens to each other, we would like the hit rate to be measured in the same way so that the intrinsic errors associated with the techniques used are consistent across the comparison. Unfortunately, given the absence of unambiguous standards for defining the outcome of a crystallisation trial, the output data from two facilities cannot be directly compared. The comparison of screens described below use only data from the C3 crystallisation laboratory, which is an ISO-certified crystallisation laboratory run by CSIRO that accepts samples from external and internal clients (Newman, 2011). All samples are set up by a limited number of operators, using standard (vapour diffusion) protocols. The screens themselves are made up within the laboratory, and are quality controlled to reduce the variation between the different instances of the blocks (Newman *et al*., 2012; Wilson *et al*., 2020). There is no restriction on the samples set up in the facility, beyond being appropriate for the robotics. Samples are spun down before being used, but there are no requirements that the samples are suitable (e.g. sufficiently pure, or sufficiently concentrated) for crystallisation. The metadata around each crystallisation plate prepared by the C3 includes details of what was set up for each plate (drop sizes, drop ratios, the screen that was used, temperature of incubation, etc) along with information about when the plate was imaged, and any scores associated with the images collected for the plate. This data is stored in a relational database which has evolved from the CM database provided in the early 2000’s from Rigaku Automation. There are two sources of image score information – human scores generated by users, and scores generated by the MARCO autoscoring algorithm (Watkins *et al*., 2019). Although tools with suggested scores (and associated colours) are provided to the users of C3 to make scoring images easy (Rosa *et al*., 2018; Rosa *et al*., 2020), there is no requirement that users use the tools, nor that they use the tools in the way that is expected. This was made abundantly clear to us in the early days of the C3 lab. One of the scores provided in the list of possible human scores is ‘Null Experiment’, which is associated with a strong magenta colour. This classification, which we assumed would be given to an image where the drop was missing or with some other, obvious pathology, was used consistently by one particular user to mark the A1 position of a plate once they had looked at it, simply because they liked the colour. They explained this only after we expressed our concern at the consistency of our problem setting up their A1 droplets! Thus the human annotations are both noisy and incomplete. Despite the limitations of the human classification, the success rates of crystallisation trials within C3 were calculated using human ‘Crystal’ scores, rather than the MARCO autoscores, for a number of reasons. Firstly, the autoscoring was implemented in 2018, and not all of the older images have been autoscored. Secondly, the MARCO autoscoring seems to over-classify images into the ‘Crystal’ class. Finally, although the human scores are noisy, the limitations of the human scoring should be independent of the screens. Figure 2 shows the initial screens offered in the C3 laboratory, the number of times they have been set up, and the (human scored) success rate. Only screens that have been set up at least 100 times are recorded on the graph. The Shotgun I screen has been set up more often than any other screen, possibly because it was seen to be successful initially, or more likely, simply because it became the default screen to use at the start of a crystallisation campaign. The Shotgun I screen and the JCSG+ screen (Newman *et al*., 2005) – which has also been set up a large number of times in the C3 - have 23 conditions in common (25% of each screen). The older Index Screen and Crystal Screen HT (called CS CSII C3 in the C3 lab), both from Hampton Research, have about 1/3 of the screening conditions in common with Shotgun I. Despite the large overlap between these screens, the outcomes from the screens are quite different. The Shotgun I screen shows a 43.7% success rate – meaning that on average, for every 100 times the shotgun screen has been set up, 44 times a human has assigned a score of ‘Crystal’ to at least one of the images collected over time from the 96 drops of the screen. This compares to 37.7% for JCSG+, another screen generated by cherry picking successful conditions from commercial screens (Page *et al*., 2003; Newman *et al*., 2005). Although the difference in success rate is only 6%, that means that Shotgun I is 16% more successful than JCSG+. Similarly, Shotgun I is almost 45% more successful than Crystal Screen HT, and 34% more successful than Index. The overall number of instances of each screen also needs to be considered – for the four screens discussed above, each has been set up in C3 at least 500 times, which is still only a small fraction of the usage of the screens over the larger structural biology community and will have sampled only a small number of possible protein/protein complex samples. Finally, the screens used in C3 are all prepared in-house, and although based on the commercially available screens, may differ sufficiently from those available commercially to alter the results that the purchased commercial screens would have produced.

**Figure 2.**
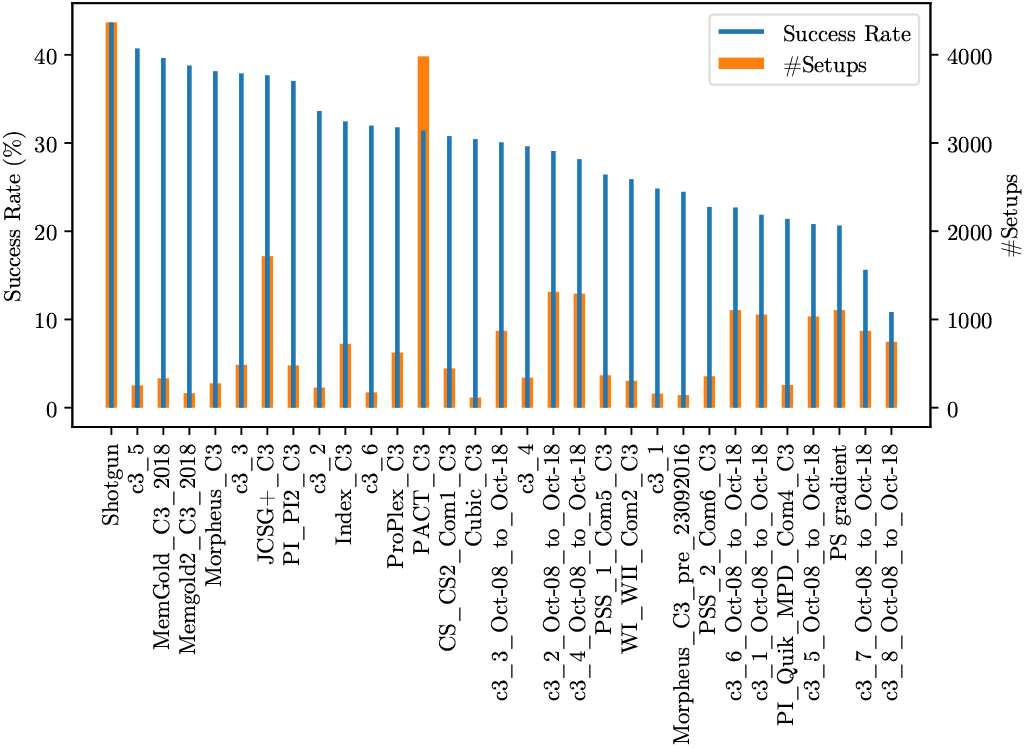
The C3 offers its users a limited number of initial screens. Many of these are based on commercial conditions. This figure shows how many times a screen has been set up in the C3 facility, (orange bars) as well as the success rate (number of times a human has scored one or more images from one or more inspections of a single screen setup/number of setups), blue bars.

Datamining the PDB to generate successful screens is not the only way of producing efficient screens. Screens can also be evolved to become better. In C3 we offer a set of eight 96-condition screens (C3 set, consisting of screens C3 1 → C3 8) which contain 768 conditions (many of them from commercial screens), but which were sorted so that each C3 x screen contained conditions with a similar major factor. See Table 1.

**Table 1.** C3 x screens and contents

| Screen Name | Contents |
| --- | --- |
| C3.1 | Peggy: Low MW (liquid PEGs), diverse buffers, pH range and salts. |
| C3.2 | Peggy: Low-Mid MW, diverse buffers, pH range and salts. |
| C3.3 | Peggy: 3350 and 4000, diverse buffers, pH range and salts. |
| C3.4 | Peggy: Mid-High MW, diverse buffers, pH range and salts. |
| C3.5 | Salty: ammonium sulfate, other sulfates, citrates |
| C3.6 | Salty: various phosphates, chlorides, acetates, formates, tartrates |
| C3.7 | Organics: MPD, small branched polymers (Jeffamines), hexanediol. |
| C3.8 | Organics: Alcohols. |

After the initial C3 x set had been in use for a decade or so, we examined the screens, and reformulated the screens so that conditions which had never produced a (human) crystal score were discarded and made up new conditions (using data on frequency and concentration of different chemicals from the PDB to guide the guesses) to replace those which had been removed. This has led to a remarkable improvement in this set of 8 screens: one screen (C3 5) improved its success rate by a factor of 2 (from 20% to 40%). The two screens C3 7 and C3 8 contain organics and alcohols and seem to be neither particularly popular amongst C3 users nor particularly successful and have not been set up 100× since reformulation, so we have no trustworthy data on how the reformulation has changed these two screens.

### 3.2. Analysis of Shotgun I outcomes

Along with measuring the overall success rate of Shotgun I, we can extend the analysis to examine the performance of individual conditions within the screen. There were a few surprises that came from this analysis. Firstly, we discovered that the code used to create the original Shotgun I had a bug which reduced all acetate salts to sodium acetate, which was thus over-represented in the screen, leading to 6 conditions containing sodium acetate whilst the corresponding commercial conditions had either ammonium acetate (4 conditions) or calcium acetate (2 conditions). Secondly, the successful commercial condition containing 2.4 M sodium malonate pH 7 (found as condition 27 of Hampton Research’s Index Screen) was accidentally transcribed as 1.4 M sodium malonate pH 7 in the original Shotgun I screen. Only two commercial screens contain 1.4 M sodium malonate pH 7, and one of those (Axygen CP-Custom-VI) is no longer available, so this condition will not have been sampled often, and is thus unsurprisingly not found often in the PDB. However, the condition is not the worst performing of the Shotgun I conditions (it ranked 77/96). This suggests that the conditions that are found multiple times in the PDB are not simply intrinsically ‘good’ but more likely are the result of some balance between being both reasonable conditions and well sampled conditions.

The Shotgun I screen conditions were ranked by the number of times the condition was found in the NR-SCC. Thus the condition in A1 of Shotgun I was the commercial condition found most often in the PDB in 2014: (0.2 M magnesium chloride, 0.1 M tris chloride pH 8.5, 30 % w/v polyethylene glycol 4000). This condition was the 12^th^ most successful according to the C3 usage; the most popular was the very similar condition C9: (0.2 M magnesium chloride, 0.1 M tris chloride pH 8.5, 25 % w/v polyethylene glycol 3350). One possible reason for this is the combination of PEG 3350 (which is known to contain PO_4_ (Wilson *et al*., 2020)) and magnesium, which could lead to crystals of magnesium phosphate, along with protein crystals. The C3 results strongly favour conditions containing polyethylene glycol; the first condition containing a salt as the primary factor (the most abundant chemical in the condition) was ranked 26^th^, and still contained some PEG: (2 M ammonium sulfate, 0.1 M sodium hepes pH 7.5, 2 % v/v polyethylene glycol 400). The least successful condition was the single condition that contained 2-methyl-2,4-pentanediol (30 % v/v 2-methyl-2,4-pentanediol, 0.1 M sodium acetate-acetic acid pH 4.6, 0.02 M calcium chloride), although the low pH of this condition could contribute to its ranking as well. All of the conditions with a final measured pH of less than 5 were found in the bottom half of the rankings, and took rankings 95 and 96. A complete listing of the rankings of the Shotgun I screen can be found in the supplementary information (Shotgun Hit rate.xlsx). This file gives the (normalised) score for both human and MARCO scores, along with the ranking from those scores.

### 3.3. Co-hits

Our analysis of the success of the individual conditions of Shotgun I from the trials of the screen in C3 showed that there were co-hits, where the same protein tended to crystallise in a consistent pattern within the screen. Examining these showed that often the subset of conditions which tend to produce similar outcomes are related. For example, Shotgun I condition B3 (0.2 M magnesium chloride; 25 % w/v polyethylene glycol 3350; 0.1 M bis-tris chloride, pH=5.5), tends to co-hit with Shotgun I condition C8 (0.2 M magnesium chloride; 25 % w/v polyethylene glycol 3350; 0.1 M bis-tris chloride, pH=6.5), these two conditions differ only by one pH unit. These two conditions were equally successful overall (B3 was given a ‘crystal’ score 186 times, C8 was scored ‘crystal’ 185 times), but that does not show that these two conditions co-hit. Co-hits occur when the *same sample* gives crystals in both conditions. Recognising the existence of co-hits does two things. Firstly, it brings home the importance of ensuring that both the raw success of a condition and its similarity to other conditions need to be considered when creating a new screen. Secondly, it provides some experimental data against which the theoretical C6 distance metric (or any other condition similarity metric) can be tested. The same observation about co-hits and condition similarity was made almost 20 years ago in the analysis by Page *et al* (Page *et al*., 2003), where they used this insight to generate a minimal core set of 67 conditions from a redundant set of 480 commercial conditions used in a Structural Genomics project.

The C6 distance metric is a heuristic designed to measure the similarity between two screens. Although what we mean by ‘similarity’ is not precise in the sense of comparing the chemistry of the conditions, it is certainly correct to say that the C6 metric should estimate conditions which are likely to crystallise the same protein as similar and conditions which are unlikely to crystallise the same protein as dissimilar. We therefore test the validity of the C6 metric by comparing it with the co-hit score. For the Shotgun I screen, we find a correlation coefficient of 0.32 between co-hits and C6 score (Fig. 3).

**Figure 3.**
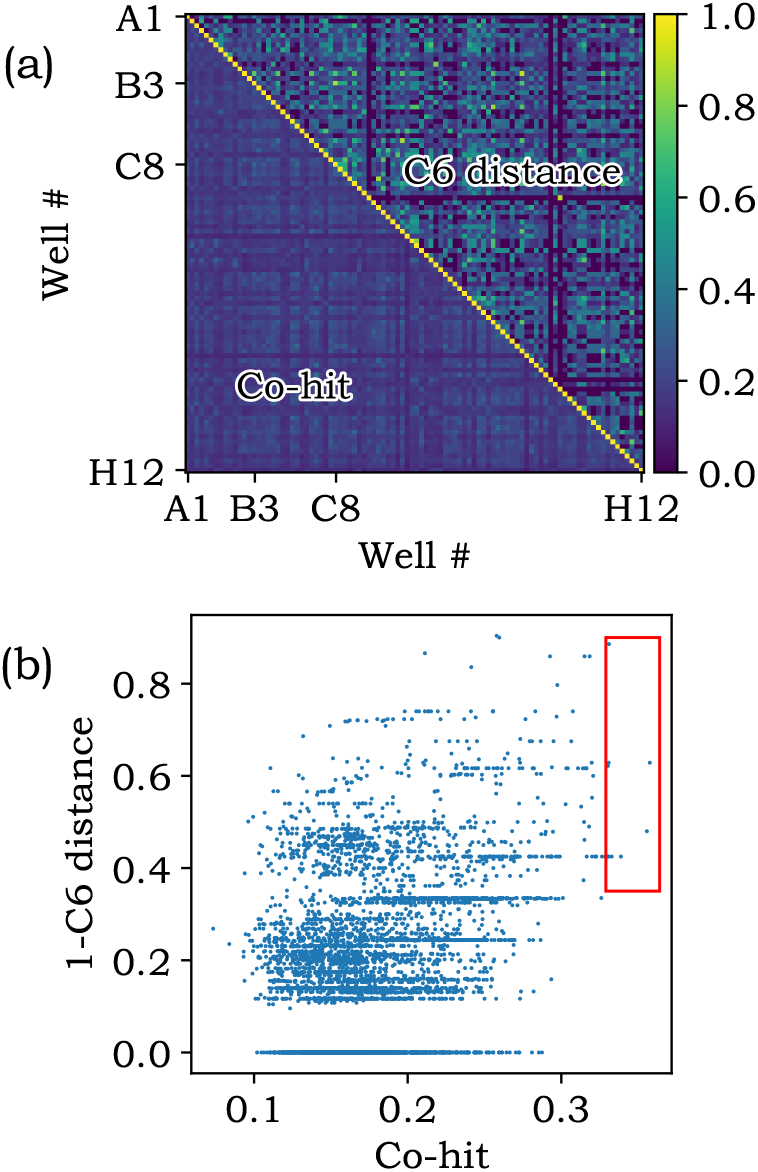
(a) The co-hit and [1 - C6] distance metric evaluated for each pair of the 96 well Shotgun I screen. All values along the diagonal are 1 in both cases. (b) Each point is a pair of conditions for which the co-hit and C6 distance have been evaluated. Identical pairs (diagonal in (a)) have been removed in (b) as they trivially increase the correlation even if C6 had no predictive power. The red box encloses the top 10 condition pairs by co-hit, which correspondingly have relatively low C6 scores.

Before calculating the correlation coefficient we remove identical conditions (for which the co-hit score is trivially 1 and C6 distance is trivially 0) as including this data biases the correlation coefficient towards 1 even if the C6 distance score were an otherwise completely random predictor. This result validates the C6 metric for the first time, demonstrating that although it is not a strong predictor, it does in fact give a better than random estimate of how likely two conditions are to crystallise a given protein given one of the conditions is known to crystallise that protein. We performed the same calculation for other screens for which data has been recorded at C3 and in every case found a positive correlation coefficient (Table. B.3).

Having a wider collection of co-hit data from the crystallisation community would enable the development of structured (*e*.*g*. Bayesian) approaches to optimisation. For example, with the co-hit data it is also possible to calculate the conditional probability that condition A will crystallise a protein given condition B crystallised it:

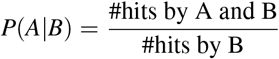

### 3.4. Parsing Problems

The parsing of the R280 fields found in the PDB is noisy. From our manual checks the current parsing tool correctly returns the crystallisation chemical factors only 80% of the time, and this assumes that the text in the R280 is error free. The more factors a condition contains the less likely it will have been transcribed into the R280 correctly, and subsequently parsed correctly. This biases our results; for example, the Morpheus screen (Gorrec, 2009) is a popular screen available from Molecular Dimensions, but it is a complicated screen, where each condition contains anywhere from 5 to 10 individual factors. Many of the Morpheus conditions are described as combinations of mixtures. The spreadsheet from Molecular Dimensions describing the screen lists the conditions in terms of mixtures (*e*.*g*. Condition A1: 0.06 M Divalents, 0.1 M Buffer System 1 6.5, 30% v/v Precipitant Mix 1), rather than as explicit chemical factors (Condition A1: 0.03 M calcium chloride; 0.03 M magnesium chloride; 0.1 M MES-imidazole, pH=6.5; 10 % w/v polyethylene glycol 20000; 20 % v/v polyethylene glycol monomethyl ether 550). As a result, we see few conditions from Morpheus in our parsed R280 data, even though the string ‘Morpheus’ is found in 513 R280s (of the 147709 PDB idents that contain the R280 field). A deeper analysis of R280 fields reveals that only 379 of the 513 R280s containing the string ‘Morpheus’ were parsed at all, and of those, only 2 were correct after parsing. There were a multitude of problems with the R280s that led to the poor results. One of the most frequent issues is that the parser is unable to distinguish chemicals in the protein formulation from chemicals in the crystallisation condition. Another frequent problem was that the R280 text was missing one or more factors, or that the factors were not enumerated individually (*e*.*g*. ‘0.03 M of both magnesium chloride and calcium chloride’). We even found one well populated R280 field that contained no information about the final crystallisation condition at all (6GJT). From a manual inspection of the 379 conditions the parser attempted to translate, the most successful Morpheus conditions found in the PDB data were conditions A2 and B2, both of which were found 11 times. A summary of the frequency of the individual conditions found in the Morpheus HT screen can be found as Supplementary Information (Morpheus counts.xlsx)

Some suggestions for how to alleviate some of the issues with parsing the PDB are presented in Appendix A.

### 3.5. NR-SCC and D-CCC in 2021

The purpose of creating the NR-SCC is to ensure that depositions in the PDB of identical proteins do not bias the rankings of the crystallisation conditions, as the same protein may have been crystallised successfully using the same conditions many times. For example, Hen Egg White Lysozyme is found hundreds of times in the PDB, often crystallised using the same condition (1 M sodium chloride, 100 mM sodium acetate pH 4.6) – so without correction, that condition would appear incredibly effective for all proteins. Any condition should gain one count only for each different protein it crystallises. We use sequence to perform the filtering as it is a relatively unique identifier of a protein. Two PDB entries are considered to be of the same protein if each chain is at least 95% identical, and the same number of chains are present. We use 95% identity rather than 100% to allow for errors in sequencing, to include tags and potentially single site mutants of the same protein. PDB entries are clustered in two steps. First, BLAST (Altschul *et al*., 1997) is used to cluster PDB chains by 95% identity, second, PDB IDs are clustered by chain cluster. These clusters are then assumed to be of identical proteins.

We use sequence similarity to ensure conditions are not double-counted in the rankings. There is however a growing body of evidence suggesting that sequence as a proxy for structure could be useful to determine likely primary crystallisation chemicals for a given protein based on its sequence (Lu *et al*., 2012; Lynch *et al*., 2020) - although our previous work (Abrahams & Newman, 2019) found no correlation between *complete* crystallisation conditions and sequence similarity.

A comparison of the pH range and the number of factors in the conditions found associated with the D-CCC and the NR-SCC shows that the distributions appear not to have altered from the distributions found in 2014 (see Appendix B).

Theoretically, any crystallisation condition could be found multiple times in the PDB, not just those from commercially available screens. However, in practise only commercial condition are found repeated in the R280. Two things are responsible. Firstly, the ubiquity of commercial screens in essentially all crystallisation laboratories. Secondly, the improbability that different groups, optimising hits from different proteins, would end up with the same chemical condition. If there were a global approach to optimisation this would be less unlikely - however a clear path from a crystallisation hit to an optimised condition is not yet available.

### 3.6. Shotgun II

The seven years since our original analysis of the PDB have resulted in an extra 50,000 X-ray structures being deposited, along with the introduction or wide adoption of new commercial screens (*e*.*g*. Morpheus and Midas from Molecular Dimensions, JBScreen LCP from Jena BioScience (Caffrey, 2015)) into the structural biology community. These new screens, if sampled often, and/or if particularly successful, could conceivably change what commercial conditions are found frequently in the PDB.

#### 3.6.1. Maximum Coverage Problem

The underlying design principle of the Shotgun I screen is that the conditions which crystallise the largest number of distinct proteins are assumed to be the most likely to crystallise unknown proteins, where ‘distinct’ is defined through the construction of the NR-SCC, detailed in Section 3.5. However, the approach taken in Shotgun I of selecting the highest ranking proteins in the NR-SCC does not in fact lead to an optimal selection of conditions. The problem of finding the conditions which crystallise the largest space of distinct proteins is an instance of the Maximum Coverage Problem (https://en.wikipedia.org/wiki/Maximum coverage problem). Given a set *S*_1_ whose elements are subsets of *S*_2_, and *k >* 0, find the subset of *S*_1_ of length *k* which is the largest subset of *S*_2_. In our case, the elements of *S*_1_ are the conditions found in the PDB, and *S*_2_ is the NR-SCC. To find the optimal solution naively, a brute force approach would need to evaluate *C*(sizeofNR −SCC, 96) ≈ 10^300^ possible screens. In fact, the maximum coverage problem is known to be NP-hard, meaning it cannot be optimally solved in polynomial time^**2**^ with any known algorithm. However, a solution can be approximately found in polynomial time using a greedy algorithm, which is known to have the best worst-case-performance of any polynomial time algorithm (Feige, 1998), achieving an approximation ratio of 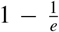, that is, a coverage of at least ≈ 0.632 times that of the the best solution (which is not necessarily 100% coverage). Other more complex algorithms may exist to find a closer approximation, however, considering that NR-SCC conditions do not overlap much (as demonstrated by only a small improvement in the greedy algorithm over selecting the highest ranking conditions) it is unlikely a significantly better selection is possible. Employing the greedy algorithm results in the 96 conditions of Shotgun II having a coverage of 5465 NR-SCC proteins, a small improvement over the 5448 found by simply selecting the top ranked conditions, as was done in 2014. The two resulting screens (Shotgun II, and Shotgun II greedy) are very similar, having 89/96 conditions in common.

#### 3.6.2. Diversity in Shotgun Screens

To optimize for the greatest possible chance of hits we considered two measures of diversity when developing the Shotgun II screen. Firstly, we considered how much of the NR-SCC would be covered by the screen. This describes ‘diversity in protein space’. The second measure was ‘diversity in chemical space’, where we used the C6 distance to set limits on how similar each pair of conditions in the screen could be. As suggested by the positive correlation between co-hits and the C6 distance metric, this is likely to be a good proxy for how likely two conditions are to crystallise the same proteins. In principle, only the protein diversity is important. However, a simple count of protein clusters as found in the NR-SCC is not a true measure of diversity. We would need to quantify the diversity of the sequence/structure space covered by the NR-SCC - the lower limit of 95% sequence identity which delineates the clusters ensures that some of the clusters are quite similar.

Cycles of internal comparison were used to create a set of six Shotgun II candidates. The first Shotgun II was produced by simply applying the greedy algorithm as described in Section 3.6.1. The remaining five screens included an increasingly stringent diversity criterion, so that the six Shotgun II candidates had pairwise minimum C6 distance thresholds of 0 → 0.5. Each condition in a Shotgun II candidate screen was compared to the other 95 conditions. If two conditions were found to be within a given C6 similarity distance then the less popular condition was discarded, and the next ranked condition was inserted. This process was repeated until convergence. See Figure 4. This process might be thought of as an attempt to remove conditions that cohit, which also might be considered as removing duplicate conditions falling into a single hotspot in chemical space. Notice that even after applying the diversity requirement the least popular condition (condition 96) in the Shotgun II-0.2 screen has 28 instances in the NR-SCC. This suggests that even if the R280s containing Morpheus conditions had been parsed correctly (section 3.4), no Morpheus condition (the best which were found 11 times) would have made it into the screen. In 2014, the least popular condition was found 20 times in the NR-SCC. Conditions found only in newer screens would have to be very successful to overcome the disadvantage they have of being found only in screens that could have been set up in recent years. This could happen if, say, crystallisation campaigns only set up newer screens, but it appears that the more recent screens are set up along with, rather than instead of, screens containing more established crystallisation conditions.

**Figure 4.**
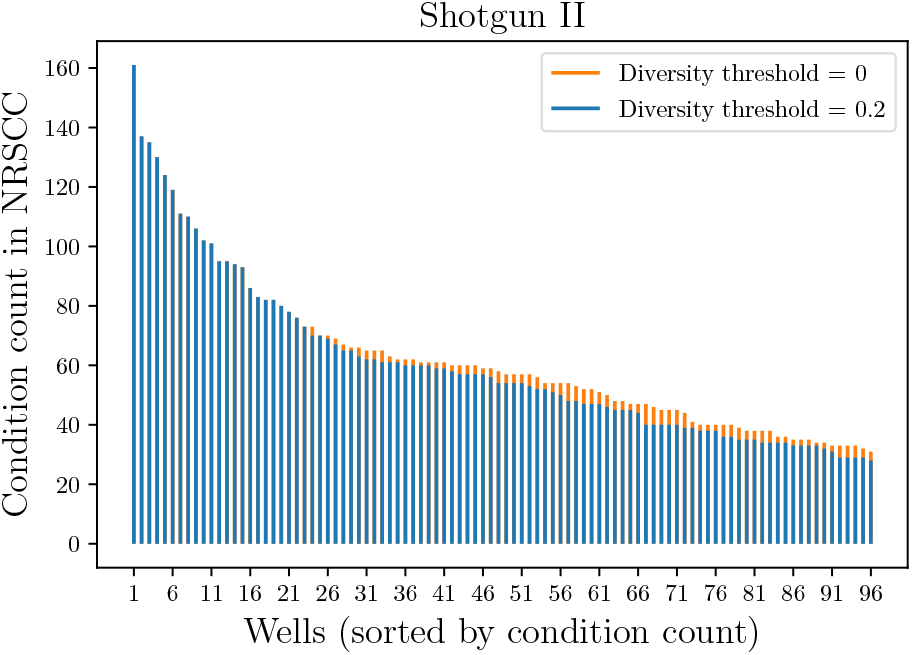
Histogram of the condition counts in the NR-SCC of the Shotgun II screen with no diversity filtering applied, and with filtering applied until every condition pair is dissimilar by at least 0.2 by the C6 metric. The highest scoring conditions are not removed by diversity filtering, hence there is no change until well 20. Beyond well 20 some conditions have been removed, however because the ranking distribution has a long flat tail the resulting screen is not composed of significantly lower ranked conditions.

There were two effects of increasing the distance between each condition in a screen. Firstly, the number of the NR-SCC clusters covered by the screen decreases. Secondly, the number of distinct chemicals in the screen increases. Notice that although the total count of NR-SCC decreases with increasing chemical diversity, different clusters are found in the more chemically diverse screens. Figure 5 and Table 3 show the effect of the increasing divergence on the total number of PDB Idents covered by the screen. For use in the C3 laboratory, we chose the screen Shotgun II-0.2 (with a C6 diversity criterion of 0.2) as our replacement for the older Shotgun I screen, as it seemed to be a good balance between NR-SCC coverage and ease of manufacture (which corresponds to the number of distinct chemical stocks required). All six Shotgun II screens are available in Supplementary information.

**Table 2.** Top ten cohits found from an analysis of the Shotgun I screen, along with the similarity [1-dissimilarity] of the two conditions from the C6 metric.

| Condition 1 | Condition 2 | Cohit probability | [1-C6] Score |
| --- | --- | --- | --- |
| ammonium sulfate 0.2M<br>polyethylene glycol 3350 25.0% w/v<br>bis-tris chloride 0.1M ph 5.5 | lithium sulfate 0.2M<br>polyethylene glycol 3350 25.0% w/v<br>bis-tris chloride 0.1M ph 5.5 | 0.358 | 0.629 |
| polyethylene glycol 3350 20.0% w/v<br>potassium sodium tartrate 0.2M | disodium tartrate 0.2M<br>polyethylene glycol 3350 20.0% w/v | 0.356 | 0.480 |
| ammonium chloride 0.2M<br>polyethylene glycol 3350 20.0% w/v | polyethylene glycol 3350 20.0% w/v<br>ammonium formate 0.2M | 0.339 | 0.425 |
| ammonium nitrate 0.2M<br>polyethylene glycol 3350 20.0% w/v | polyethylene glycol 3350 20.0% w/v<br>ammonium iodide 0.2M | 0.333 | 0.425 |
| magnesium chloride 0.2M<br>polyethylene glycol 3350 25.0% w/v<br>bis-tris chloride 0.1M ph 5.5 | magnesium chloride 0.2M<br>polyethylene glycol 3350 25.0% w/v<br>bis-tris chloride 0.1M ph 6.5 | 0.331 | 0.886 |
| lithium sulfate 0.2M<br>polyethylene glycol 3350 25.0% w/v<br>bis-tris chloride 0.1M ph 6.5 | ammonium sulfate 0.2M<br>polyethylene glycol 3350 25.0% w/v<br>bis-tris chloride 0.1M ph 6.5 | 0.331 | 0.629 |
| polyethylene glycol 3350 20.0% w/v<br>sodium malonate 0.2M ph 7.0 | disodium tartrate 0.2M<br>polyethylene glycol 3350 20.0% w/v | 0.331 | 0.425 |
| sodium acetate 0.2M<br>sodium cacodylate 0.1M ph 6.5<br>polyethylene glycol 8000 18.0% w/v | sodium mes 0.1M ph 6.0<br>polyethylene glycol 8000 20.0% w/v<br>sodium acetate 0.2M | 0.330 | 0.622 |
| magnesium formate 0.2M<br>polyethylene glycol 3350 20.0% w/v | magnesium chloride 0.2M<br>polyethylene glycol 3350 20.0% w/v | 0.330 | 0.425 |
| magnesium formate 0.2M<br>polyethylene glycol 3350 20.0% w/v | polyethylene glycol 3350 20.0% w/v<br>magnesium acetate 0.2M | 0.329 | 0.425 |

**Table 3.** Comparing the Shotgun I screen with the different Shotgun II screens

| Shotgun II- | NR-SCC Coverage | Distinct Chemicals | Stocks Needed | Shotgun I Overlap | Internal Diversity |
| --- | --- | --- | --- | --- | --- |
| 0.0 | 5465 | 51 | 61 | 66 | 0.917 |
| 0.1 | 5438 | 51 | 61 | 63 | 0.921 |
| 0.2 | 5310 | 52 | 62 | 65 | 0.926 |
| 0.3 | 4708 | 64 | 75 | 51 | 0.951 |
| 0.4 | 4106 | 68 | 79 | 43 | 0.954 |
| 0.5 | 3340 | 81 | 94 | 32 | 0.945 |

**Figure 5.**
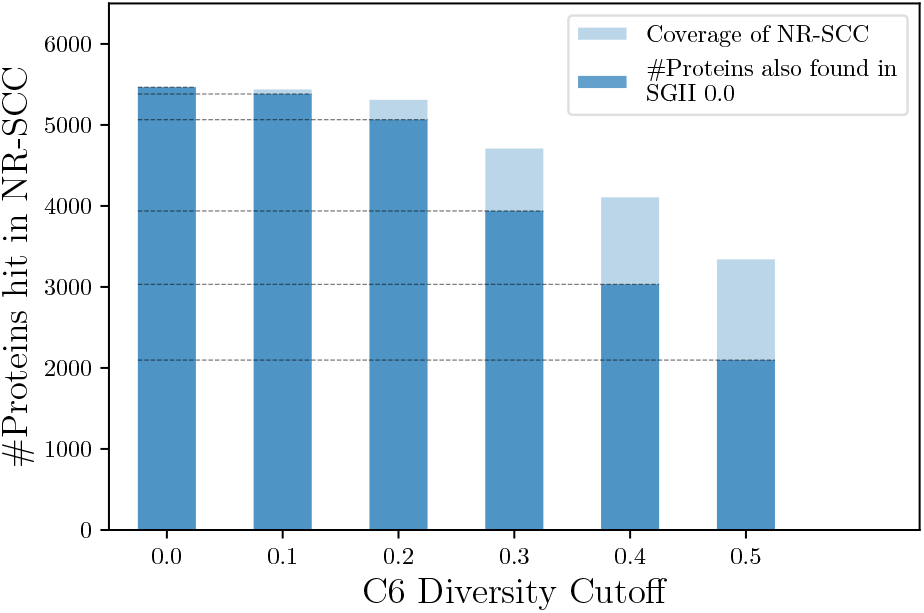
Plot of crystals (‘Hits’) versus Internal Diversity Cutoff. The Hits axis gives the total count of structures from the NR-PDB which have crystals that grew from the conditions in the different versions of the Shotgun II screen (light blue). In dark blue are the number of conditions in that screen which are also in the 0.0 screen (for 0.0 there is total overlap). As the internal diversity (average C6 distance) becomes greater, the screens become more diverse, that is, they cover more of chemical space, but become less effective in recapitulating crystals used to produce PDB structures. This is seen in the decreasing fraction of shared proteins between 0.0 and 0.X screens: the proteins hit by the 0.5 screen include less than half of those hit by the 0.0 screen, accounting for slightly more than half of the 0.5 screen itself. The Shotgun II screen with a 0.2 internal diversity cutoff was chosen to maximise both Hits and chemical diversity.

## 4. Conclusion

Revisiting the Shotgun analysis from 2014 has been valuable for a number of reasons. Firstly, this analysis allows us to compare the success rate of different screens given data from C3 on both the rate of setup and rate of forming crystallisation hits, which in turn validates data-mining approaches to find hotspots in screening space. Further, this allows us to extend the analysis both in terms of the number of data points going into the analysis, but also by including the concept of diversity when building the new screen. Finally, this study gives some insight into the use of the commercial screens within the structural biology community, suggesting that new screens are often used in addition to the more traditional screens, rather than replacing them.

## Acknowledgements

Thanks to Tom Peat, Alex Caputo and Albert Ardevol Grau for discussions, and the users of the C3 facility for providing the samples on which this work is based.

## Appendix A Parsable PDB data

The quality of the data collected in the R280 field has not noticeably improved in the 7 years since this analysis was last performed. Using the same scripts as in 2014, Figure A.1 shows that although there are proportionally slightly more parsable entries in 2021, the rate of successful parsing (which is an indication of the quality of the data in the R280 field) hasn’t changed much.

The crystallisation condition that produced the crystal(s) used in a diffraction experiment tends to be either a commercial condition unchanged, or more likely, an optimised condition based on one or more commercial conditions. During the structure deposition, we suggest that the depositor be given the option of selecting one of those two options, and then be given a list of commercial conditions from which to select either the successful condition or the set of conditions upon which the final condition was based. The R280 field would be used only to clarify the primary information. There should be a clearly documented and separate place for describing the protein formulation.

**Figure A.1.**
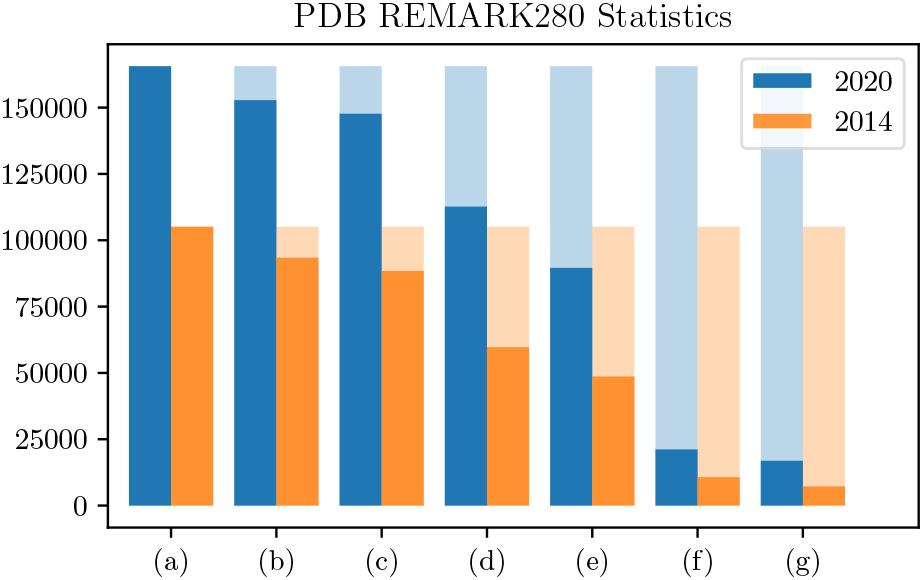
Histogram of the process of generating the data used to generate the Shotgun II screen (in blue) and the Shotgun I screen (in orange). (a) Total number of depositions in the PDB. (b) Number of X-Ray depositions. (c) Number of depositions with a REMARK280. (d) Number of depositions with a non-empty R280 that can be parsed. (e) Estimated number of correctly parsed depositions. (f) Number of depositions with crystallisation condition found in a commercial screen. (g) Number of NR-SCC conditions found in a commercial screen. The a table of the values used to generate the image can be found in sup Fig(x). The analyses follow the same trend, showing that there has been no increase in the proportion of parsable R280 field over time, nor any increase in the quality (*i*.*e*. ability to be correctly parsed) of the field. However, the latest analysis does provide twice as many data points for the comparison with the commercial screening conditions.

**Figure A.2.**
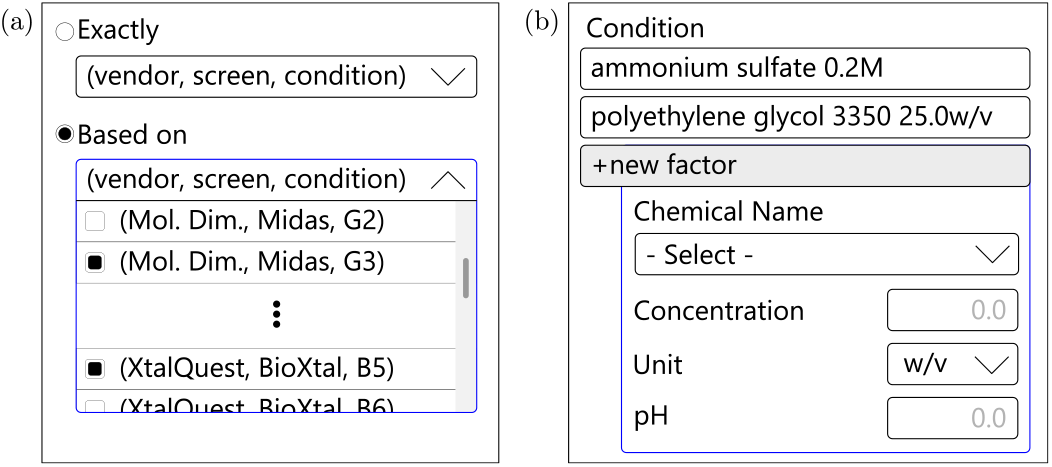
Along with the current free text R280 it would be very useful to have a formatted source of crystallisation data. Approximately 20% of X-ray structures deposited in the PDB have been determined from crystals obtained from commercial screens directly. Many of the other crystals have been grown from conditions based on results from commercial screening. A form pre-filled with commercial conditions could be used to capture these cases. (a) Shows a suggestion for a simple interface to pick either a single or multiple conditions from commercial screens. (b) Shows an interface where a user can access a library of known chemicals to build a unique condition. The current R280 would be then used to add detail - *e*.*g*. Base condition was supplemented with 10% of Hampton Research Additive Screen HT condition B9.

## Appendix B Comparisons of NR-SCC and D-CCC in 2021

The pH ranges of both the conditions found in the NR-SCC and the D-CCC are similar, as they were in 2014. As well, the average number of factors in each condition in the NR-SCC is a little higher than found in the D-CCC, again replicating what was seen in 2014. As in 2014, in 2021 the chemical found most often in the NR-SCC is polyethylene glycol 3350, and as before, the lists of the the most commonly found chemicals in the NR-SCC and the D-CCC don’t match well.

**Figure B.1.**
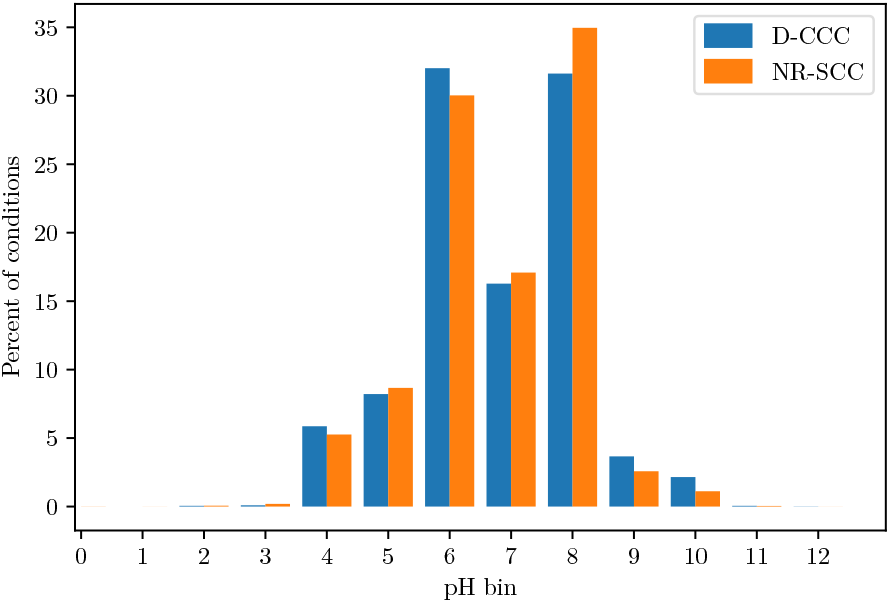
Histogram of pH

**Figure B.2.**
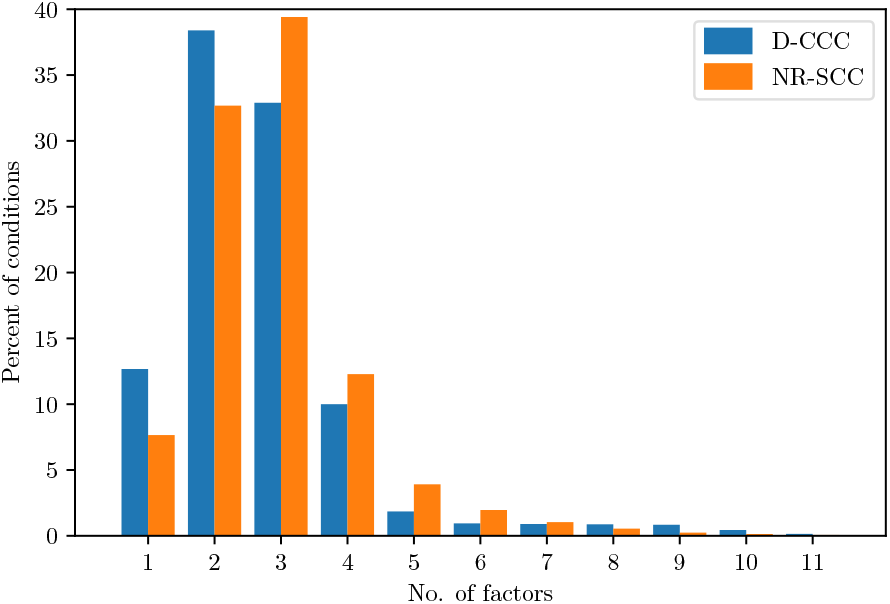
Histogram of factors

**Table B.1.** Top 10 chemicals in the NR-SCC and D-CCC

| NR-SCC Chemical Name | Count | D-CCC Chemical Name | Count |
| --- | --- | --- | --- |
| polyethylene glycol 3350 | 19320 | polyethylene glycol 4000 | 1207 |
| ammonium sulfate | 14063 | sodium chloride | 1199 |
| sodium chloride | 10124 | ammonium sulfate | 1010 |
| polyethylene glycol 4000 | 9692 | polyethylene glycol 400 | 935 |
| polyethylene glycol 8000 | 7424 | magnesium chloride | 789 |
| magnesium chloride | 7305 | polyethylene glycol 3350 | 783 |
| glycerol | 6320 | polyethylene glycol 8000 | 755 |
| lithium sulfate | 4217 | 2-methyl-2,4-pentanediol | 738 |
| polyethylene glycol 400 | 4151 | glycerol | 587 |
| polyethylene glycol 6000 | 3769 | lithium sulfate | 546 |

**Table B.2.** Top 10 buffers in the NR-SCC and D-CCC

| NR-SCC Buffer Name | Count | D-CCC Buffer name | Count |
| --- | --- | --- | --- |
| tris chloride | 11531 | sodium hepes | 1438 |
| sodium hepes | 11307 | tris chloride | 1390 |
| sodium mes | 6410 | sodium mes | 1007 |
| sodium acetate | 5704 | sodium acetate | 951 |
| bis-tris chloride | 4467 | trisodium citrate | 709 |
| trisodium citrate | 3962 | sodium cacodylate | 483 |
| sodium cacodylate | 3141 | bis-tris chloride | 328 |
| bis-tris propane chloride | 1711 | bis-tris propane chloride | 310 |
| imidazole | 1205 | imidazole | 218 |
| sodium bicine | 779 | sodium bicine | 188 |

**Table B.3.**
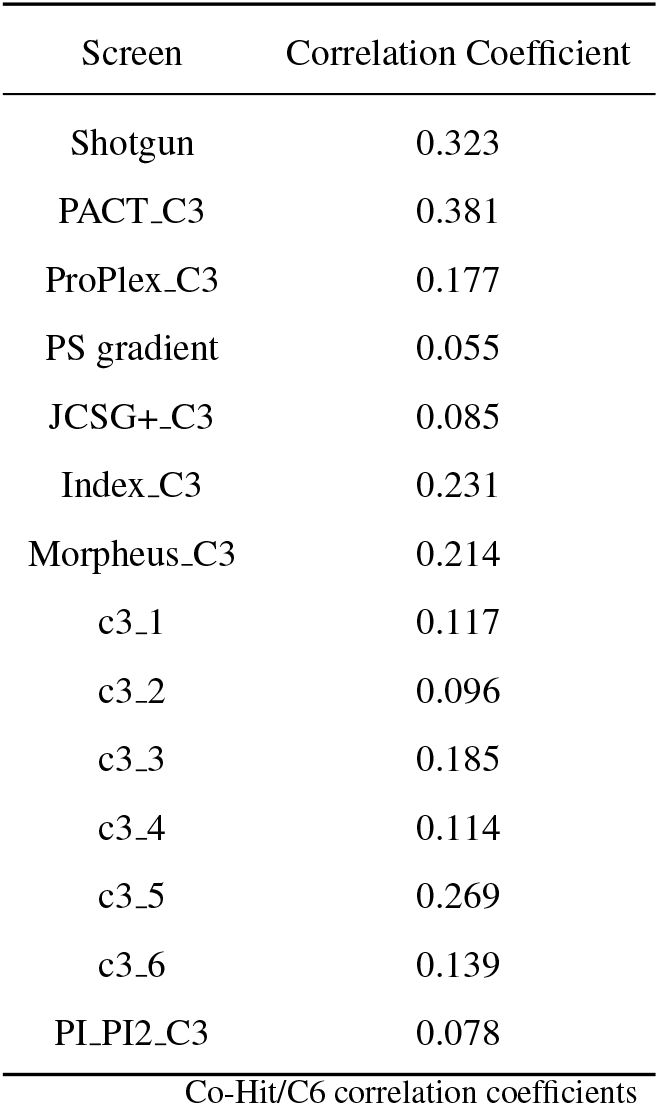

## Footnotes

1 The size of the space can be estimated using *C(5000,3)*, assuming 1000 different chemicals, which can be sampled at 5 different concentrations or pHs, and 3 factors in each condition.

2 ‘Time’ refers to how the number of computer operations scales with the problem size (here the number of clusters in the NR-SCC)

## Notes

### Competing Interest Statement

The authors have declared no competing interest.

